# ADAD2 regulates heterochromatin in meiotic and post-meiotic male germ cells via translation of MDC1

**DOI:** 10.1101/2021.07.29.454317

**Authors:** Lauren G. Chukrallah, Aditi Badrinath, Gabrielle G. Vittor, Elizabeth M. Snyder

**Affiliations:** Department of Animal Science, Rutgers University, New Brunswick

**Keywords:** Meiosis, mRNA translation, Germ cells, RNA binding proteins, Chromatin, Spermatogenesis

## Abstract

Male germ cells establish a unique heterochromatin domain, the XY-body, early in meiosis. How this domain is maintained through the end of meiosis and into post-meiotic germ cell differentiation is poorly understood. ADAD2 is a late meiotic male germ cell specific RNA binding protein, loss of which leads to post-meiotic germ cell defects. Analysis of ribosome association in *Adad2* mutants revealed defective translation of *Mdc1*, a key regulator of XY-body formation, late in meiosis. As a result, *Adad2* mutants show normal establishment but failed maintenance of the XY-body. XY-body defects are concurrent with abnormal autosomal heterochromatin and ultimately lead to severely perturbed post-meiotic germ cell heterochromatin and cell death. These findings highlight the requirement of ADAD2 for *Mdc1* translation, the role of MDC1 in maintaining meiotic male germ cell heterochromatin, and the importance of late meiotic heterochromatin for normal post-meiotic germ cell differentiation.

**Summary:** Chukrallah et al. demonstrate ADAD2 is required for normal meiotic heterochromatin in male germ cells and loss leads to post-meiotic cell death defining ADAD2 as a heterochromatin maintenance factor.

## Introduction

Male germ cell differentiation is accompanied by radical changes in the chromatin landscape as cells progress from mitosis to meiosis to spermiogenesis (Kota and Feil, 2010). Of particular importance is the formation early in meiosis of the XY-body, a heterochromatic region encapsulating the X and Y chromosome. Failure to form an XY-body results in meiotic cell arrest and infertility (Ichijima et al., 2011, Abe et al., 2020). Thus, while factors initiating this event have been well described, how the XY-body’s unique chromatin state is maintained through the later stages of meiosis and whether this is important for post-meiotic steps of male germ cell differentiation is unknown.

In early meiotic spermatocytes, pericentric heterochromatin (PCH) of both sex chromosomes and autosomes is composed of similar epigenetic marks (Magaraki et al., 2017). However, with the onset of chromosome synapsis, PCH and other heterochromatin marks diverge between the sex chromosomes and the autosomes resulting in two unique chromatin compartments, autosomal PCH and the XY-body, composed of sex chromosome PCH and other sex-chromosome enriched epigenetic marks (Namekawa et al., 2006). These compartments ultimately give rise to distinct compartments in the post-meiotic germ cell, with autosomal PCH forming a region termed the chromocenter (CC) and sex chromosome PCH forming post-meiotic sex chromatin (PMSC) (Namekawa et al., 2006, Haaf and Ward, 1995). While genetic evidence demonstrates XY-body formation is required in early and mid-meiosis to suppress synapsis checkpoints and silence sex chromosome gene expression (meiotic sex chromosome inactivation – MSCI) (McKee and Handel, 1993), it is unknown whether stringent maintenance in late meiosis is necessary for post-meiotic germ cell differentiation and how maintenance may be achieved.

Two key mediators of XY-body formation are Breast Cancer 1 (BRCA1) and mediator of DNA-damage checkpoint protein 1 (MDC1). In male germ cells, BRCA1 is required for the initial establishment of X chromosome γH2AX through its recruitment of ATR and TOPBP1, ultimately forming X chromosome PCH (Broering et al., 2014). MDC1, on the other hand, interacts with γH2AX (Ichijima et al., 2011, Savic et al., 2009) and leads to spreading of the γH2AX mark throughout the X and Y, giving rise to the XY-body (Ichijima et al., 2011, Namekawa et al., 2006). Although the MDC1-γH2AX pathway has been traditionally associated with DNA damage response (DDR) (Savic et al., 2009, Blanco-Rodriguez, 2012), a growing consensus contends that meiotic male germ cells leverage it to facilitate chromatin-induced silencing of the X and Y chromosome early in meiosis. In support of this, MDC1 either directly or indirectly influences the XY-body localization of multiple chromatin remodeling proteins required to establish the unique epigenetic signature of the XY-body (Ichijima et al., 2011). MDC1 and BRCA1 remain high throughout meiosis (Ahmed et al., 2007, Turner et al., 2004) suggesting they may be important in the maintenance of the XY-body, however mutation of either results in germ cell loss prior to meiotic completion (Ahmed et al., 2007), thus their roles in late meiosis and post-meiotic events are unclear.

Several lines of evidence suggest the chromatin state of the XY-body late in meiosis is important for post-meiotic germ cell development. The chromatin composition of the CC and PMSC closely mimics their meiotic counterparts. Further, mutation of the MDC1-interacting protein Scm polycomb group protein like 2 (SCML2), leads to dysregulation of spermatid chromatin (Luo et al., 2015). Mutation of a second meiotic chromatin remodeling protein, bromodomain testis associated (BRDT), alters heterochromatin in both meiotic compartments during late meiosis and displays distinct CC abnormalities coupled to defects in sperm maturation (Berkovits and Wolgemuth, 2011). As spermatid chromatin state is thought to establish the nuclear topology necessary for genome compaction in the sperm head (Haaf and Ward, 1995, Meyer-Ficca et al., 1998), these observations couple meiotic chromatin state with final germ cell genome remodeling. The underlying mechanisms driving this phenomenon are, as yet, unknown.

Translation regulation, mediated by RNA binding proteins (RBPs), is a hallmark of meiotic and post-meiotic male germ cells (Braun et al., 1989, Kleene et al., 1984). As a result, there is a solid understanding of how RBPs facilitate translation repression during meiosis (Yang et al., 2007). While there has been no equivalent study of how RBPs enhance translation during meiosis and the physiological importance thereof, genetic models have shown that abnormal translation initiation can lead to failed meiotic progression (Sun et al., 2010). Further, translation activation during meiosis is dynamically regulated in both mammalian oocytes (Susor et al., 2016) and yeast (Brar et al., 2012). Coupled to this, male germ cells face the additional complexity arising from MSCI which prevents expression of X chromosome genes, such as *Scml2*, late in meiosis (Mueller et al., 2008) where they are required for meiotic events (Hasegawa et al., 2015). Together, these observations suggest positive regulation of translation may be an important regulator of male germ cell meiosis however the relevant RBPs, their mRNA targets, and the physiological outcome of abnormal translation have yet to be elucidated.

ADAD2 is a germ cell specific RBP necessary for post-meiotic germ cell differentiation as *Adad2* mutant germ cells fail to undergo the final stage of development, spermiogenesis (Snyder et al., 2020). However, the molecular underpinnings of germ cell loss in the *Adad2* mutant are unknown. Paradoxically, ADAD2 is expressed and detected exclusively in mid- and late meiotic germ cells suggesting it regulates processes in meiosis important for post-meiotic events (Snyder et al., 2020). One of the key events of post-meiotic differentiation is the establishment of a compacted haploid genome via the sequential replacement of histones first with transition proteins followed by protamines (Braun, 2001, Yu et al., 2000, Zhao et al., 2004). However, it remains unclear whether epigenetic programming in the meiotic cell directly influences post-meiotic chromatin transitions.

Here we demonstrate ADAD2 is required for normal translation of *Mdc1* late in meiosis. In *Adad2* mutants, defective *Mdc1* translation gives rise to aberrant heterochromatin in both autosomes and the sex chromosomes of late meiotic spermatocytes. These defects are retained in haploid spermatids which ultimately undergo arrest and apoptosis. Our studies define the mechanism of germ cell death in *Adad2* mutants and highlight the central role of MDC1 in maintaining heterochromatin in both chromatin compartments of the late meiotic germ cell, the importance of this maintenance for normal post-meiotic germ cell chromatin, and the key role of translation regulation therein.

## Results

### *Adad2* mutant round spermatids exhibit abnormal chromatin structure

*Adad2* mutants (*Adad2^em3^*, referred to herein as *Adad2^M/M^*) display post-meiotic germ cell loss (Snyder et al., 2020). In our effort to understand the molecular underpinnings of the *Adad2^M/M^* phenotype, we examined post-meiotic germ cell morphology in *Adad2^M/M^* testis. The chromocenter, a morphological hallmark of the post-meiotic round spermatids, is a heterochromatic region central to spermatid nuclear organization (Berkovits and Wolgemuth, 2011) and can be easily assessed via DAPI staining. As expected, this analysis (Fig. 1A) showed wildtype round spermatids nearly always contained a single, well defined chromocenter. In contrast, *Adad2^M/M^* round spermatids contained multiple regions of dense DAPI staining, often displaying dramatic increases in chromocenter-like foci relative to wildtype. We confirmed the heterochromatic nature of these morphological features via immunofluorescent staining with the heterochromatin mark HP1α (Fig. 1B).

**Figure 1.**
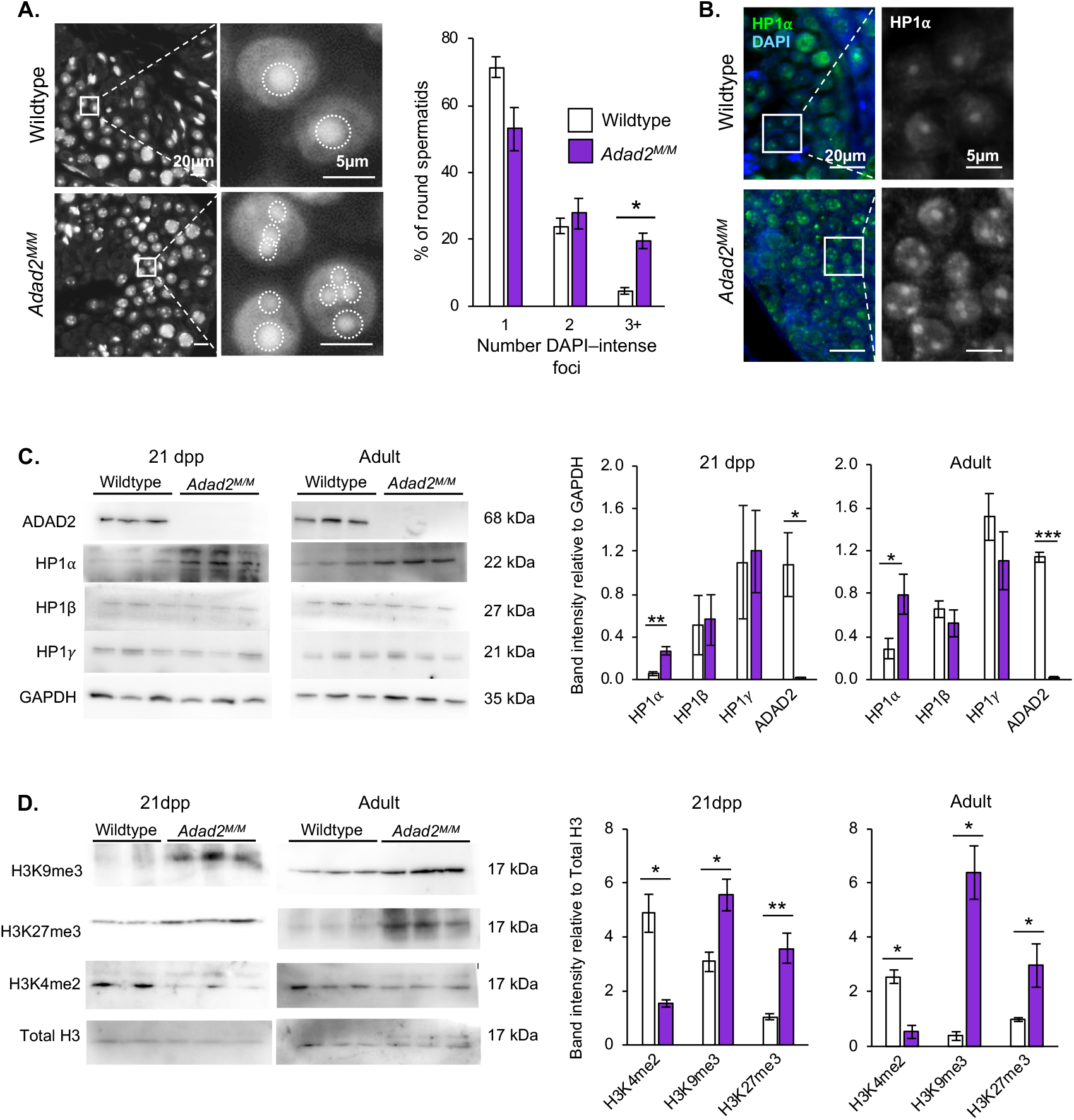
*Adad2* mutant meiotic spermatocytes and post-meiotic spermatids have abnormal localization and increased levels of heterochromatin. **A)** DAPI-stained adult wildtype and *Adad2* mutant (*Adad2^M/M^*) testis cross sections show a single DAPI-intense focus (chromocenter) in wildtype spermatids and an increased number of DAPI-intense foci in mutant spermatids. Left panel - 20x magnification. Right panel - 63x. Dashed circles - DAPI-intense foci. Quantification of DAPI-intense foci demonstrates a significant increase in mutant spermatids. **B)** Immunofluorescence of HP1α in wildtype and *Adad2* 21 dpp testis sections reveals a single HP1α foci in wildtype spermatids and multiple foci in mutant spermatids consistent with abnormal chromocenter number in mutants. Left panel - 40x magnification, green - HP1α, blue - DAPI. Right panel - 63x, HP1α only. **C)** Western blot of HP1 proteins in 21 dpp and adult whole testis lysate from wildtype and *Adad2^M/M^* samples (biological triplicate / genotype) with ADAD2 and GAPDH as genotype and loading controls, respectively. Quantification of band intensity confirms a significant increase of HP1α in mutants at both ages. **D)** Western blot of select epigenetic marks in 21 dpp and adult whole testis histone lysate from wildtype and *Adad2^M/M^* samples (21 dpp wildtype biological duplicate; biological triplicate for all else) with total histone 3 (H3) as a loading control. Quantification of band intensity shows increased heterochromatin, measured by H3K9me3 and H3K27me3, and decreased euchromatin, measured by H3K4me2. Throughout, error bars represent one standard deviation and significance was calculated using Student’s *t*-test.

The increase in chromocenter number observed in *Adad2^M/M^* may be a result of abnormal heterochromatin condensation or an overall heterochromatin increase. To distinguish between these two possibilities, we quantified the abundance of three chromobox (CBX) proteins (referred to herein as HP1 proteins: HP1α, HP1β, and HP1γ), well-characterized markers of heterochromatin with distinct expression and localization patterns in meiotic and post-meiotic germ cells (Charaka et al., 2020, Chevillard et al., 1993, Brown et al., 2010) (Fig. 1B). We examined both adult and 21 dpp testes, which are dominated by post-meiotic and late meiotic germ cells respectively, to estimate when heterochromatin changes may arise in the mutant. Of the three HP1 proteins, only HP1α was found to be significantly increased, with this increase most apparent at 21 dpp (Fig. 1C). While HP1β and HP1γ are expressed early in the pachytene phase of meiosis, HP1α functions much later in meiosis (Takada et al., 2011). Thus, the increase of HP1α but not HP1β or HP1γ suggests increased heterochromatin in the *Adad2* mutants arises late in pachytene, timing that coincides with expression of ADAD2 in wildtype spermatocytes.

Although HP1α interacts with a wide range of DNA-associated proteins, it shows special affinity for K9-methylated histone H3 (H3K9me3) (Bannister et al., 2001). Thus, we examined H3K9me3 along with other epigenetic marks representative of heterochromatin (H3K27me3) and euchromatin (H3K4me2) by Western blot of 21 dpp and adult testis histone lysate (Fig. 1D). At 21 dpp*, Adad2^M/M^* testes exhibited increased levels of both H3K9me3 and H3K27me3 along with reduced levels of H3K4me2, consistent with the hypothesis that *Adad2^M/M^* spermatocytes house increased heterochromatin with a subsequent reduction in euchromatin. These differences persisted to adulthood demonstrating altered chromatin state was maintained in mutant germ cells beyond pachytene spermatocytes. In total, these findings suggest that mutant round spermatids may have an abnormal chromatin state, which could be causative to their loss in the *Adad2* mutant, and this abnormal state arises in late meiotic spermatocytes.

### *Adad2* mutation alters transcript abundance and ribosome association of specific transcripts

ADAD2 protein is exclusive to mid- and late-spermatocytes and mutants display heterochromatin defects late in meiosis suggesting the mutant phenotype is due to events in mid-to late meiosis. To better define causes leading to the heterochromatin defect in *Adad2* mutants, we performed RNA-sequencing of total RNA from 21 dpp wildtype and mutant testes which are enriched for late meiotic germ cells. This analysis (Fig. 2A) identified a moderate number of differentially expressed (DE) genes (Supp. Table 1), with the majority having reduced abundance in the mutant testes (61%). Previous work examining *Adad2^M/M^* 25dpp testes demonstrated a robust reduction in transcript abundance, most likely due to post-meiotic germ cell loss (Snyder et al., 2020). To determine if this was the case at 21 dpp as well, expression of DE genes in wildtype testicular somatic and germ cells was examined (SF 1A). This analysis showed DE transcripts reduced in the mutant were primarily expressed in wildtype meiotic and post-meiotic germ cells. Concurrently, DE transcripts increased in the mutant generally had high expression in mitotic wildtype cells, a pattern consistent with reduced numbers of meiotic and post-meiotic germ cells in the mutant. However, detailed morphological quantification of meiotic and post-meiotic cell populations (SF 1B-C) showed no cell loss in the mutant until the round spermatid stage. These findings were further confirmed by quantifying TUNEL-positive spermatocytes and spermatids in mutant adults (SF 1D-F), analyses that showed increased apoptotic germ cells were limited to post-meiotic spermatids.

**Figure 2.**
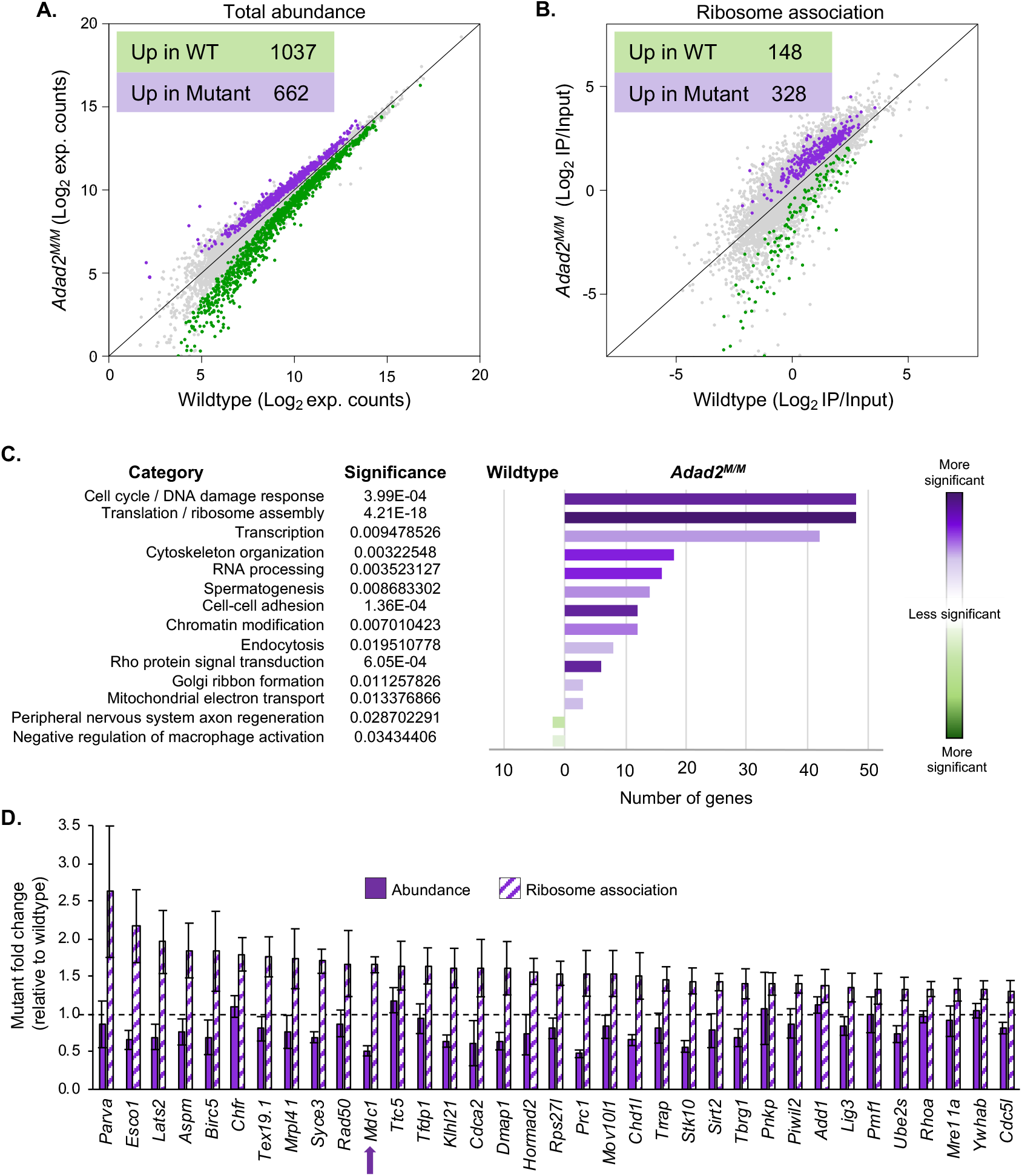
Mutation of *Adad2* results in altered transcript abundance and ribosome association of specific transcript classes. Comparison of **A)** total transcript abundance and **B)** ribosome association in wildtype and *Adad2* mutant (*Adad2^M/M^*) Ribotag samples (green - upregulated in wildtype, purple - upregulated in mutant) identifies many transcripts with differential ribosome association (DRA) in *Adad2* mutant testes. **C)** Significantly enriched ontological categories for DRA transcripts (wildtype - green and *Adad2^M/M^* – purple) identifies cell cycle and DNA damage response (DDR) as significantly enriched in transcripts with increased ribosome association in the mutant. **D)** Total abundance and ribosome association (derived from RNA-sequencing) relative to wildtype of select cell cycle and DDR related genes identified as DRA confirms increased ribosome association and minimal abundance changes in mutant testes. Dashed line represents wildtype average. Arrow indicates gene of interest, *Mdc1.* Error bars represent one standard deviation.

Toward defining the mechanisms giving rise to the *Adad2* mutant phenotype, we examined the functions of differentially expressed genes via ontological analysis (SF 1G). It should however be noted that, of the differentially expressed genes, only 4.28% exhibited a fold-change of two-fold or higher (Supp. Table 1) demonstrating abundance changes in the mutant are relatively few in number and also of low magnitude. Ontology analysis demonstrated genes associated with spermiogenesis and cell projection were significantly enriched in the wildtype testes consistent with a loss of post-meiotic germ cells in the mutant. Contrary to expectation, no dramatic changes in transcripts encoding chromatin remodeling proteins or histones were observed. Together, these observations demonstrate ADAD2 loss results in a slight reduction of meiotic germ cell transcript abundance in the absence of appreciable meiotic germ cell loss and transcript abundance changes alone cannot explain the observed heterochromatin defects.

Germ cell RNA granules are known sites of translational regulation (Lehtiniemi and Kotaja, 2018) and given the previously reported granular localization of ADAD2 in meiotic germ cells (Snyder et al., 2020) along with its relatively limited impacts on transcript abundance, we hypothesized ADAD2 may play a role in post-transcriptional regulation. To determine the impact of *Adad2* mutation on translation, we combined our *Adad2* mutant allele with the RiboTag model (Sanz et al., 2009), which expresses a cell-specific HA-tagged large ribosomal subunit protein when driven by a cell-specific Cre. In our model, RiboTag expression was driven in differentiating germ cells via *Stra8-iCre* (*RiboTag+:Stra8-iCre+*) (SF 2A). Thus, RNA-sequencing of total RNA (input) and HA immunoprecipitated (IP) ribosome-RNA complexes from wildtype and mutant *RiboTag+:Stra8-iCre+* 21 dpp testes was used to quantify ribosome association across wildtype and *Adad2* mutant differentiating germ cell transcriptomes (SF 2B). Ribosome association (RA) was expressed as a ratio of IP over input abundance (IP/Input) to correct for any genotype-driven differential transcript abundances. From this value, differential ribosome association (DRA) was determined (Fig. 2B and listed in Supp. Table 2) and most commonly showed increased ribosome association in the mutant as compared to the wildtype (69% of transcripts). Analysis of total abundance for all DRA transcripts (SF 2C) confirmed this was not due to overall reduced transcript abundance in the mutant testis. Further, transcripts with higher RA in mutants showed the highest expression in wildtype meiotic germ cells (SF 2D). Together, these results suggest *Adad2* mutation results in increased ribosome association of meiotic germ cell transcripts, in agreement with the spermatocyte-specific expression of the ADAD2 protein.

We next performed ontological analysis of DRA transcripts to determine whether differential ribosome association may lead to the *Adad2^M/M^* phenotype (Fig. 2C). While wildtype-increased DRA transcripts showed poor ontological enrichment, mutant-increased DRA transcripts had striking enrichment for translation and cell cycle functions, specifically meiosis and DNA damage response, and included many of the most profoundly impacted DRA genes (Supp. Table 2). It has been well described that meiotic male germ cells leverage the DNA damage response (DDR) pathway to establish the correct epigenetic state during meiosis (Namekawa et al., 2006), thus we focused on defining the exact impact of differential ribosome association on select DDR proteins.

### Increased ribosome association results in altered DDR protein abundance

Combined analysis of total abundance and ribosome association (Fig. 2D) showed cell-cycle and DDR transcripts had a general trend of decreased abundance and increased ribosome association in *Adad2* mutants. While increased ribosome association is assumed to be indicative of increased protein production, in some cases increased ribosome occupancy can lead to reduced protein production and subsequent transcript degradation (Brandman et al., 2012, D’Orazio et al., 2019). Thus, to test how ribosome association influenced protein abundance in the *Adad2* mutant, we examined two biologically relevant transcripts: one with only increased expression in the *Adad2* mutant (*Brca1*) and another with increased ribosome association in conjunction with moderately reduced transcript abundance (*Mdc1*). Both influence DNA damage response (Lou et al., 2003, Campbell et al., 2010, Stewart et al., 2003) and meiotic germ cell heterochromatin state, in particular XY-body formation and MSCI (Ichijima et al., 2011, Becherel et al., 2013, Royo et al., 2013) making them potential candidates for mediating heterochromatin defects in *Adad2* mutants. Western blot analysis in 21 dpp wildtype and mutant testes showed a reciprocal pattern of protein abundance, with MDC1 greatly reduced and BRCA1 dramatically increased (Fig. 3A). While BRCA1 displayed the anticipated response to increased expression, the reduction of MDC1 was unexpected. Thus, we selected several additional transcripts with abundance and ribosome association profiles similar to *Mdc1* and examined their protein abundances (SF 3A-C). In all cases, increased ribosome association in conjunction with moderately decreased transcript resulted in dramatically reduced protein abundance to a degree that could not be attributed to transcript abundance decreases.

**Figure 3:**
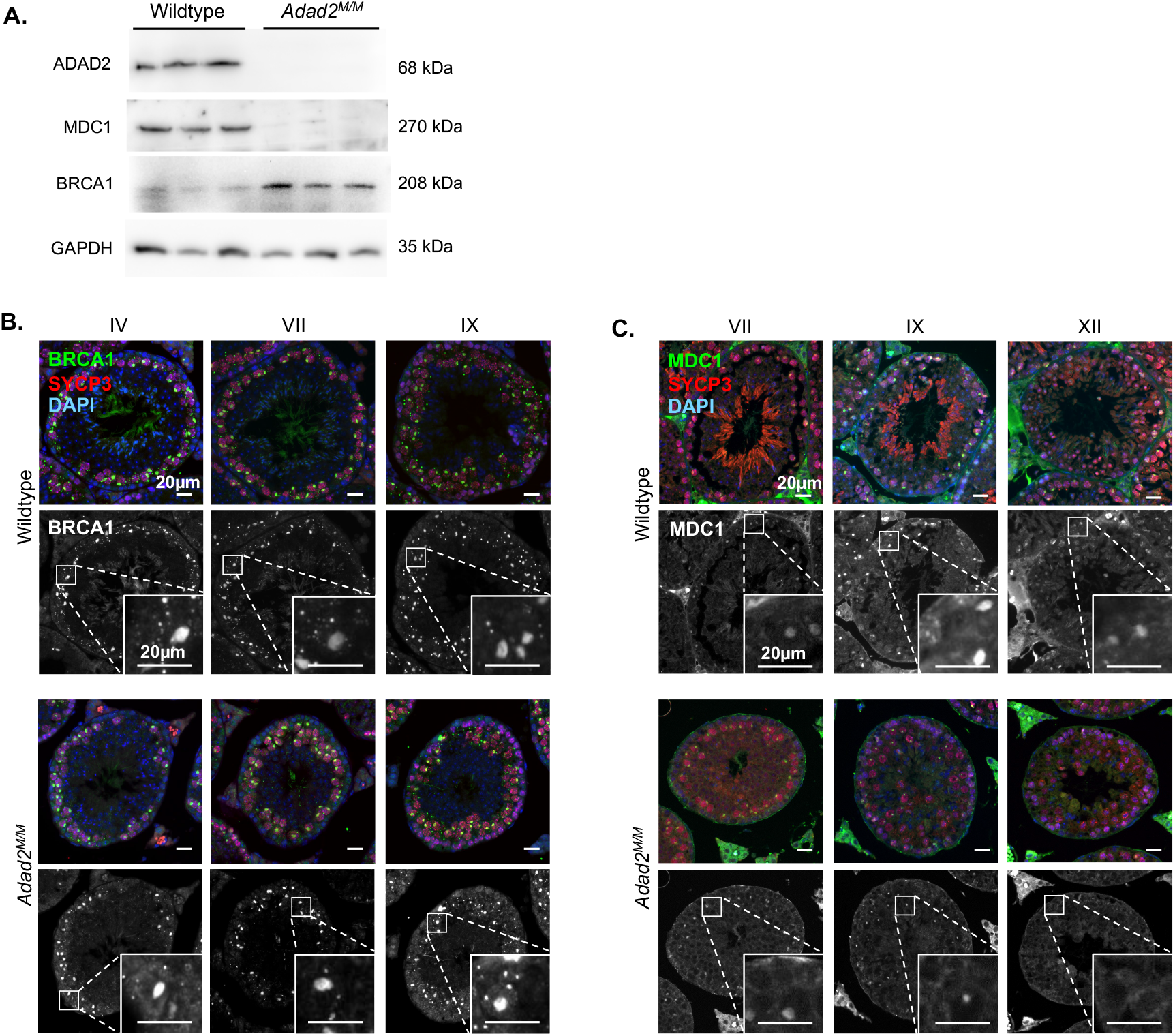
Loss of ADAD2 results in altered abundance of two key DNA damage response proteins. **A)** Western blot of MDC1 and BRCA1 in wildtype and *Adad2^M/M^* adult testis lysate shows reciprocal impacts on protein abundance in *Adad2* mutants. GAPDH and ADAD2 blots shown as loading and genetic controls (biological triplicate / genotype. **B)** Immunofluorescence of BRCA1 in adult wildtype and *Adad2^M/M^* testis sections by stage (indicated by Roman numerals) showing increase of XY-body associated BRCA1 in mutant mid-to late-stage (VII and IX) spermatocytes relative to wildtype. Color - overlays of BRCA1 (green), SYCP3 (red), and DAPI (blue). Single channel – BRCA1 alone. For all large panels - 20x magnification; insets - 40x. **C)** Immunofluorescence of MDC1 in adult wildtype and *Adad2^M/M^* testis sections by stage showing loss of MC1 in late meiotic spermatocytes (stages IX and XII). Color - overlays of MDC1 (green), SYCP3 (red), and DAPI (blue). Single channel - MDC1 alone.

Together, these results suggest ADAD2 differentially influences the abundance of two targets (BRCA1 and MDC1) important for heterochromatin remodeling in meiotic male germ cells and identify BRCA1 and/or MDC1 as potential mediators of the *Adad2* mutant phenotype. Given both are localized to and closely associated with the formation of the XY-body (Ichijima et al., 2011, Broering et al., 2014, Kogo et al., 2012), we first asked whether loss of ADAD2 and the subsequent changes in protein abundance altered their localization (Fig. 3B and C). For both, we observed normal XY-body localization and intensity in early pachytene spermatocytes. However, while BRCA1 showed increased XY-body association in both mid- and late-stage *Adad2^M/M^* spermatocytes, which normally express high levels of granule-localized ADAD2, a dramatic reduction of MDC1 was observed exclusively in late-stage spermatocytes. These observations demonstrate ADAD2 is required to maintain normal BRCA1 and MDC1 protein levels, particularly in late-stage spermatocytes, which influences their relative concentration in the XY-body.

### ADAD2 loss leads to accumulation of γH2AX on autosomes

BRCA1 and MDC1 both function in the XY-body to establish a sex-chromosome wide γH2AX domain (Broering et al., 2014, Ichijima et al., 2011)). To assess the impact of elevated BRCA1 in conjunction with reduced MDC1, we first examined total γH2AX in 21 dpp whole testes (SF 4A) which showed an overall increase. Next, we examined the localization of γH2AX in wildtype and *Adad2* spermatocytes and found that *Adad2^M/M^* spermatocytes had qualitatively normal levels of γH2AX within the XY-body throughout meiosis suggesting increased BRCA1 did not result in an expansion of the γH2AX domain. However, late pachytene and diplotene *Adad2^M/M^* spermatocytes did show significantly increased γH2AX along the axes of the autosomes relative to wildtype, a phenomenon not observed in early and mid-pachytene spermatocytes of either genotype (Fig. 4A). In order to determine whether this aberrant autosomal localization was a result of persistent DNA damage we quantified RPA2 foci, which mark sites of DNA damage for recombination (Ichijima et al., 2011, Raderschall et al., 1999), in wildtype and *Adad2* spermatocytes and found no significant difference (SF 4B & C) demonstrating normal DNA damage repair kinetics in mutant spermatocytes and suggesting the aberrant γH2AX signal is not a result of persistent or recurring DNA damage.

**Figure 4:**
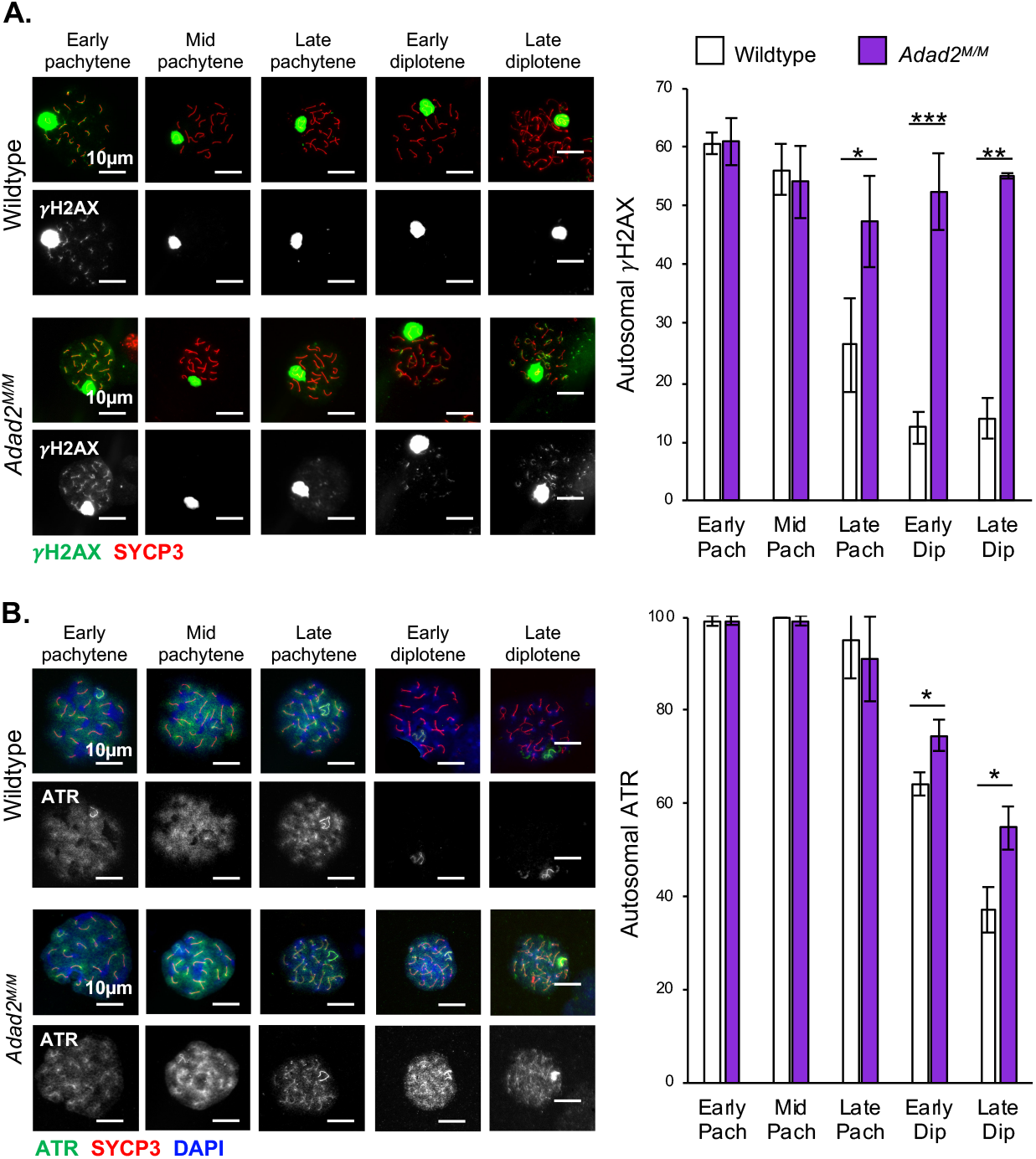
Loss of ADAD2 results in abnormal autosomal yH2AX and ATR late in meiosis. **A)** γH2AX localization and quantification by spermatocyte stage in 30dpp wildtype and *Adad2^M/M^* germ cells (40x magnification, green - γH2AX, red - SYCP3) showing mutant-specific retention of autosomal yH2AX in late pachytene through diplotene. **B)** ATR localization and quantification by spermatocyte stage in 30dpp wildtype and *Adad2^M/M^* germ cells (40x magnification, green – ATR, red – SYCP3, blue - DAPI) showing increased retention of autosomal ATR in mutant spermatocytes late in meiosis. Error bars represent one standard deviation. Significance calculated using Student’s *t*-test. For all quantifications: * p < 0.05, ** p < 0.001, *** p < 0.0001

ATR is initially recruited to the XY-body by BRCA1 early in meiosis where it phosphorylates H2AX to form γH2AX (Turner et al., 2004). γH2AX, in turn, is bound by MDC1 (Stewart et al., 2003) (Ichijima et al., 2011), which then tethers ATR to the sex chromosomes and facilitates the spread of γH2AX throughout the remainder of the sex chromosomes (Alavattam et al., 2016, Ichijima et al., 2011). To determine whether autosomal γH2AX in *Adad2* mutant spermatocytes was a function of mis-localized ATR, we quantified autosomal ATR in *Adad2^M/M^* spermatocytes. While this analysis demonstrated normal XY-body ATR in mutant spermatocytes, they also showed a persistent ATR autosomal signal through to late diplotene, a pattern not observed in the wildtype (Fig 4B). This indicates autosomal γH2AX was likely a function of ATR localization. Given normal XY-body ATR and γH2AX, we conclude increased BRCA1 had minimal impact on their behavior. In contrast, spreading of ATR to the autosomes implies loss of MDC1 leads to distinct failure of XY-body γH2AX and ATR maintenance and suggesting that the primary *Adad2* phenotype may be driven by MDC1 loss.

### *Adad2* mutant meiotic germ cells display characteristics indicative of MDC1 loss

While MDC1 is not the only ADAD2-impacted protein, it is a key modulator of epigenetic reprogramming in meiotic male germ cells. Thus, we wondered whether *Adad2* mutant spermatocytes displayed molecular phenotypes indicative of MDC1 loss. We first asked whether well characterized downstream targets of MDC1 were altered in *Adad2* mutant spermatocytes, with an emphasis on late pachytene spermatocytes. In this cell population, SCML2 localizes to the XY-body in an MDC1-dependent manner where it recruits the deubiquitinating enzyme USP7. As a result, in wildtype late pachytene spermatocytes, histone 2A lysine 119 ubiquitination (H2AK119Ub, referred to here as K119Ub) is partially excluded from the XY-body (Fig. 5A) (Luo et al., 2015, Hasegawa et al., 2015). Hence, mutation of *Scml2* or *Mdc1* results in XY-body accumulation or inclusion of K119Ub, respectively (Luo et al., 2015, Adams et al., 2018). Given the reduction of MDC1 in *Adad2* mutant spermatocytes, we assessed the localization of SCML2 activity in pachytene and diplotene spermatocytes by first examining the localization of USP7 (Fig. 5B). In wildtype spermatocytes USP7 transitions from both autosomal and XY-body localized to XY-body enriched by the end of meiosis. In contrast, while mutant spermatocytes established the correct USP7 XY-body enrichment in early and mid-pachytene spermatocytes, USP7 shifts out of the XY-body in late pachytene and diplotene spermatocytes, concurrent with an accumulation of autosomal USP7. We next asked whether this USP7 shift resulted in abnormal deposition of the epigenetic mark K119Ub (Fig. 5B). As expected based on the USP7 localization, K119Ub localization was similar between wildtype and mutant in early and mid-pachytene spermatocytes but exclusion was significantly reduced in mutant late pachytene and diplotene spermatocytes (Fig. 5C). Together, these observations demonstrate a dramatic redistribution of one MDC1-regulated pathway, including the epigenetic mark resulting from it, in *Adad2* mutant spermatocytes at the time of MDC1 reduction. This suggests that adequate MDC1 levels may be necessary to maintain proper localization of downstream networks once established.

**Figure 5:**
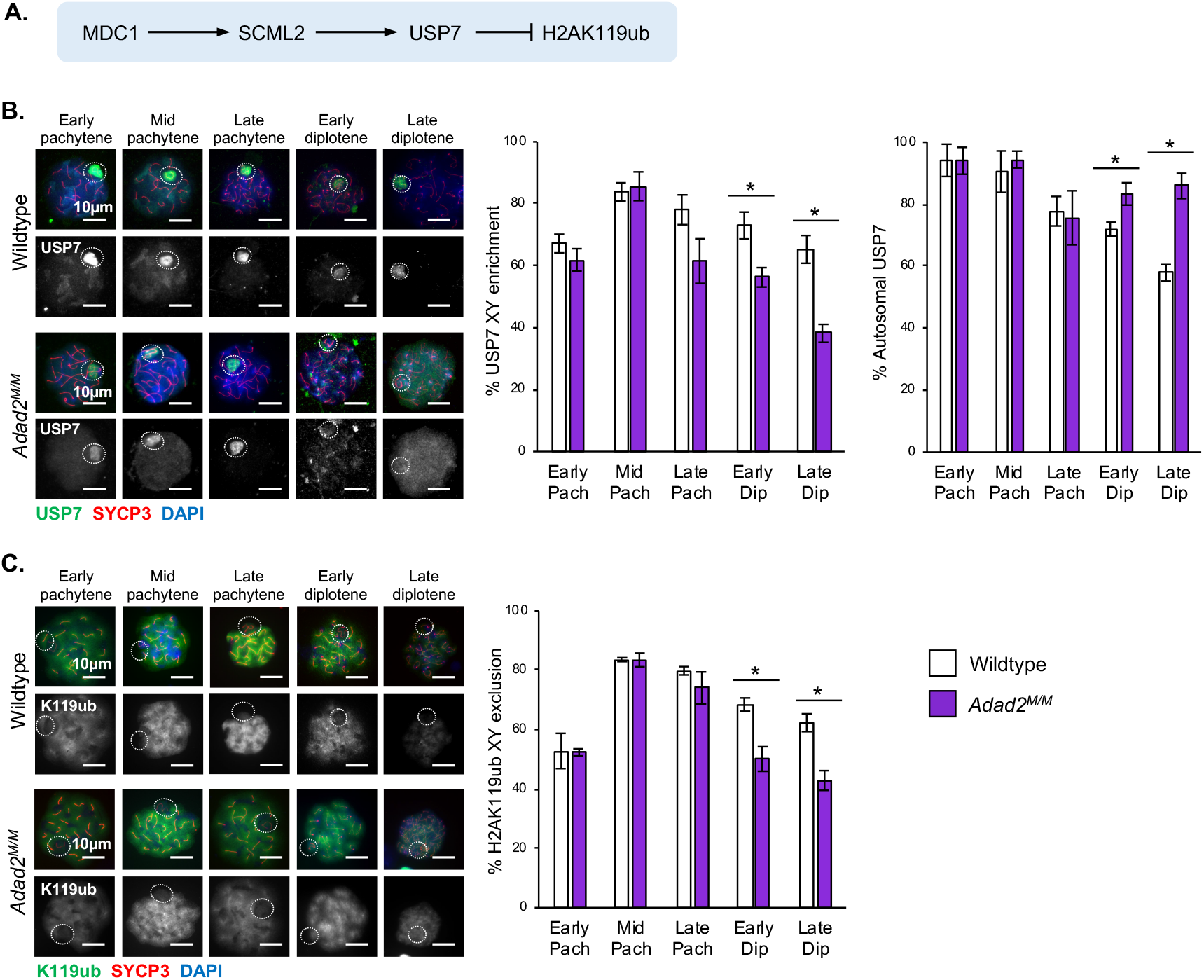
The MDC1-regulated SCML2 network shifts from the XY-body to the autosome in late-pachytene mutant spermatocytes. **A)** MDC1 regulation of USP7 and H2AK119ub via SCML2. **B)** Immunocytochemistry of USP7 and localization quantification by spermatocyte stage in 30dpp wildtype and *Adad2^M/M^* spermatocyte spreads (green - USP7, red - SYCP3, blue – DAPI) showing decreased XY enrichment and increase autosomal signal of USP7. **C)** Immunocytochemistry of H2AK119ub and localization quantification by stage in 30dpp wildtype and *Adad2^M/M^* spermatocyte spreads. (green - H2AK119ub, red - SYCP3, blue - DAPI) showing loss of H2AK119Ub XY-body exclusion in mutant late meiotic spermatocytes. Dashed circles – region containing X and Y chromosome. Error bars represent one standard deviation. Significance calculated using Student’s *t*-test. * p < 0.05, ** p < 0.001, *** p < 0.0001

In spite of the redistribution of USP7 in the mutant spermatocytes, it remained a possibility that the observed changes resulted from alterations in the expression or ribosome association of known epigenetic mark regulators. To determine if this was the case, we examined key elements of each pathway using our RNA and RiboTag-sequence analyses (SF 3D). Of these, no significant changes in abundance or ribosome association were observed in *Adad2^M/M^* samples, indicating altered epigenetic marks cannot be attributed exclusively to expression or ribosome association changes.

### MSCI is moderately defective in *Adad2* mutants

In early pachytene spermatocytes, MDC1 mediates silencing of sex chromosome expression (MSCI) via deposition or exclusion of specific epigenetic marks specifically on the sex chromosomes (Abe et al., 2020, Ichijima et al., 2011). Given the observed changes to the sex chromosome epigenetic composition, we examined MSCI in *Adad2* mutants by comparing differential expression (DE) across the autosome and sex chromosomes in 21 dpp wildtype and mutant testes (SF 5A-B). These analyses demonstrated the X chromosome was not overrepresented in the DE gene list nor was X chromosome gene expression significantly increased as a whole. This is in contrast to other models lacking MSCI in which many X chromosome genes are dramatically up-regulated (Turner, 2007). However, in spite of what appeared to be qualitatively normal MSCI in mutant spermatocytes, X chromosome DE genes were much more likely to be up-regulated as compared to autosomal DE genes (SF 5C).

While the direct influence of MDC1 on MSCI in late pachytene spermatocytes cannot be assessed due to the *Mdc1* mutant phenotype, mutation of the MDC1 target *Scml2* does result in abnormal MSCI late in meiosis (Luo et al., 2015). Thus, we compared X chromosome genes up-regulated in *Adad2* mutants with those in *Scml2* mutants and found an appreciable overlap (SF 5D). Together, these findings suggest MSCI is established normally in *Adad2* mutants but may be maintained improperly, presumably due to insufficient MDC1 protein levels leading to abnormal SCML2 function.

### Deposition of the activating mark H3K4me2 is abnormally regulated in *Adad2* mutants

While the vast majority of differentially expressed X chromosome genes in the *Adad2* mutant showed increased abundance, a select number demonstrated a decrease. Detailed analysis of these revealed half (SF 7E) belong to the relatively small number of X chromosome encoded transcripts that undergo activation after the completion of meiosis (Ernst et al., 2019). Reduced expression of these transcripts suggested there may be defects in post-meiotic gene activation of the X-chromosome, which is driven in part by deposition of the activating mark H3K4me2 (Adams et al., 2018). Supporting this notion, our initial heterochromatin characterization (Fig. 1D) demonstrated reduced H3K4me2 in mutant whole testis lysate. To further explore this, we characterized H3K4me2 XY-body deposition during meiosis, which is dependent on the MDC1-interacting protein RNF8 (Adams et al., 2018). Early in meiosis H3K4me2 is excluded from the XY-body (de Vries et al., 2012, Federici et al., 2015), a finding recapitulated in early and mid-pachytene wildtype and mutant spermatocytes. However, while H3K4me2 was enriched in the XY-body of wildtype late-stage pachytene spermatocytes, virtually no XY-body associated H3K4me2 was observed in mutant late-stage pachytene spermatocytes (Fig. 6). As for other epigenetic marks, we examined gene expression and ribosome association for H3K4 methylation genes (SF 3D) and found no significant changes. Together, these analyses suggest ADAD2 indirectly influences H3K4 methylation via its influence on MDC1, in agreement with the reported role of MDC1 in enhancing H3K4me2 in the XY-body. Further, they confirm *Adad2* mutant spermatocytes display molecular phenotypes indicative of MDC1 loss in late pachytene spermatocytes.

**Figure 6:**
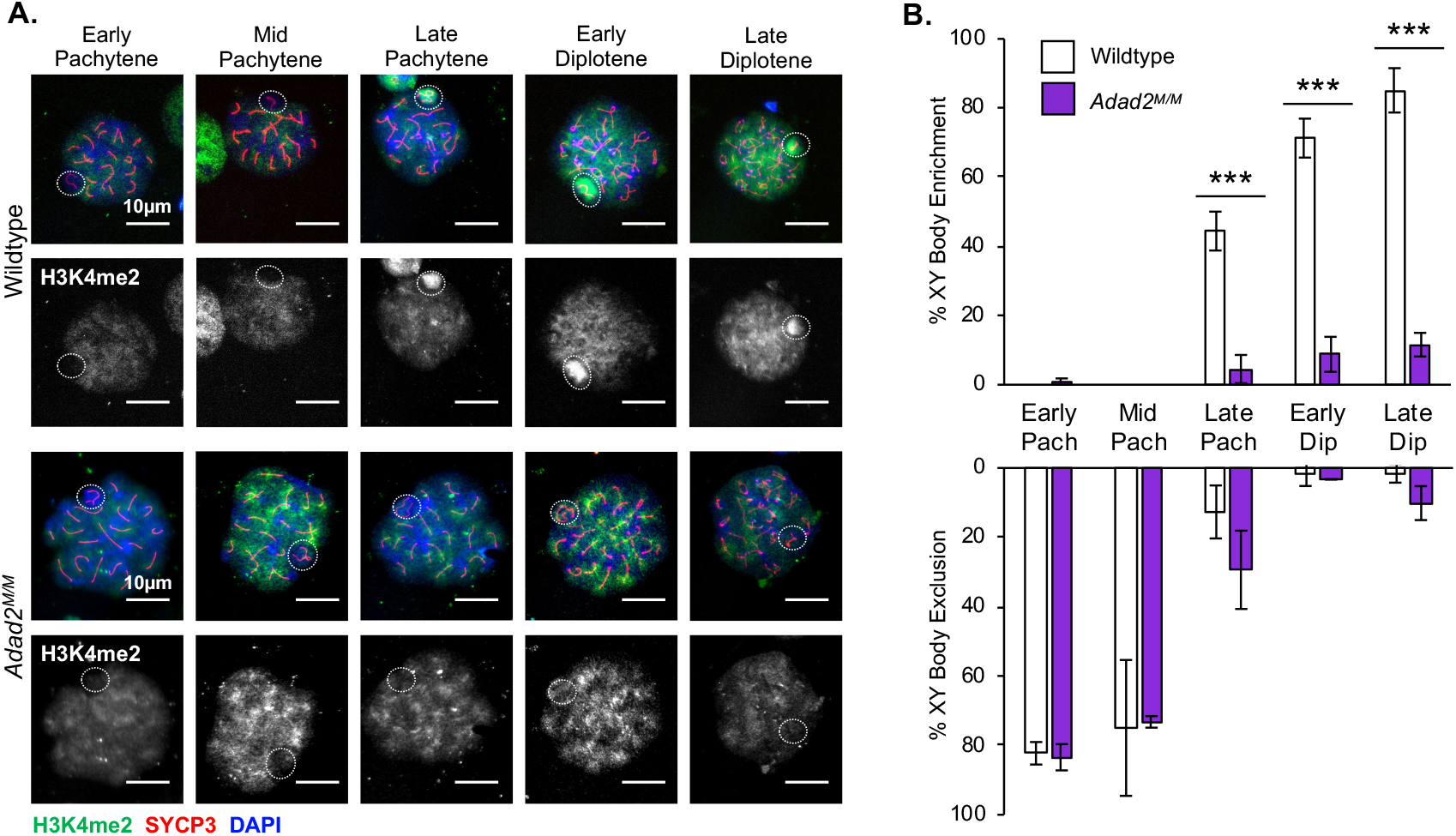
Mutant spermatocytes fail to localize H3K4me2 to the XY-body late in meiosis. **A)** H3K4me2 immunocytochemistry (green – H3K4me2, red – SYCP3, blue – DAPI, dashed circles - region containing X and Y chromosome) and **B)** quantification of localization in 30 dpp wildtype and *Adad2^M/M^* spermatocytes showing lack of XY-body accumulation in mutant spermatocytes late in meiosis. Error bars represent one standard deviation. Significance calculated using Student’s *t*-test. * p < 0.05, ** p < 0.001, *** p < 0.0001

### *Adad2* mutant spermatids have abnormalities in post-meiotic chromatin and chromatin remodeling

The heterochromatin of post-meiotic spermatids is divided into the chromocenter and a second highly heterochromatic region termed post-meiotic sex chromatin (PMSC) that contains the condensed X and Y chromosomes (Moretti et al., 2016). Given dramatic alterations in meiotic chromatin of *Adad2* mutants and the previously observed abnormalities in chromocenter morphology, we examined epigenetic marks in the heterochromatin of *Adad2* mutant round spermatids.

Each round spermatid heterochromatin region is marked by distinct combinations of histone modifications, with the chromocenter containing both H3K9me3 and H3K27me3 while PMSC is strongly H3K9me3 positive but depleted of H3K27me3 (Ichijima et al., 2012, Turner et al., 2006). Using co-immunofluorescence of H3K9me3 and H3K27me3 in *Adad2^M/M^* round spermatids (Fig. 7A), we observed the fragmented chromocenters of *Adad2^M/M^* round spermatids were marked by both H3K9me3 and H3K27me3, similar to single chromocenters in the wildtype. Further, mutant round spermatids did not display either the weakly stained DAPI or H3K9me3-only regions indicative of PMSC, suggesting *Adad2^M/M^* spermatocytes lack a distinct PMSC domain. To confirm, we performed coimmunofluorescence of H3K4me2 and H3K27me3 in *Adad2^M/M^* round spermatids (Fig. 7B) with the expectation that regions of PMSC should be strongly H3K4me2 enriched and chromocenters H3K4me2 depleted (Tatehana et al., 2020). This pattern was observed in wildtype round spermatids, however in the *Adad2* mutant spermatids H3K4me2 did not localize to any of the chromocenter-like structures and there were relatively few regions of distinct H3K4me2 enrichment or exclusion demonstrating *Adad2* mutant spermatids lack distinguishable PMSC.

**Figure 7:**
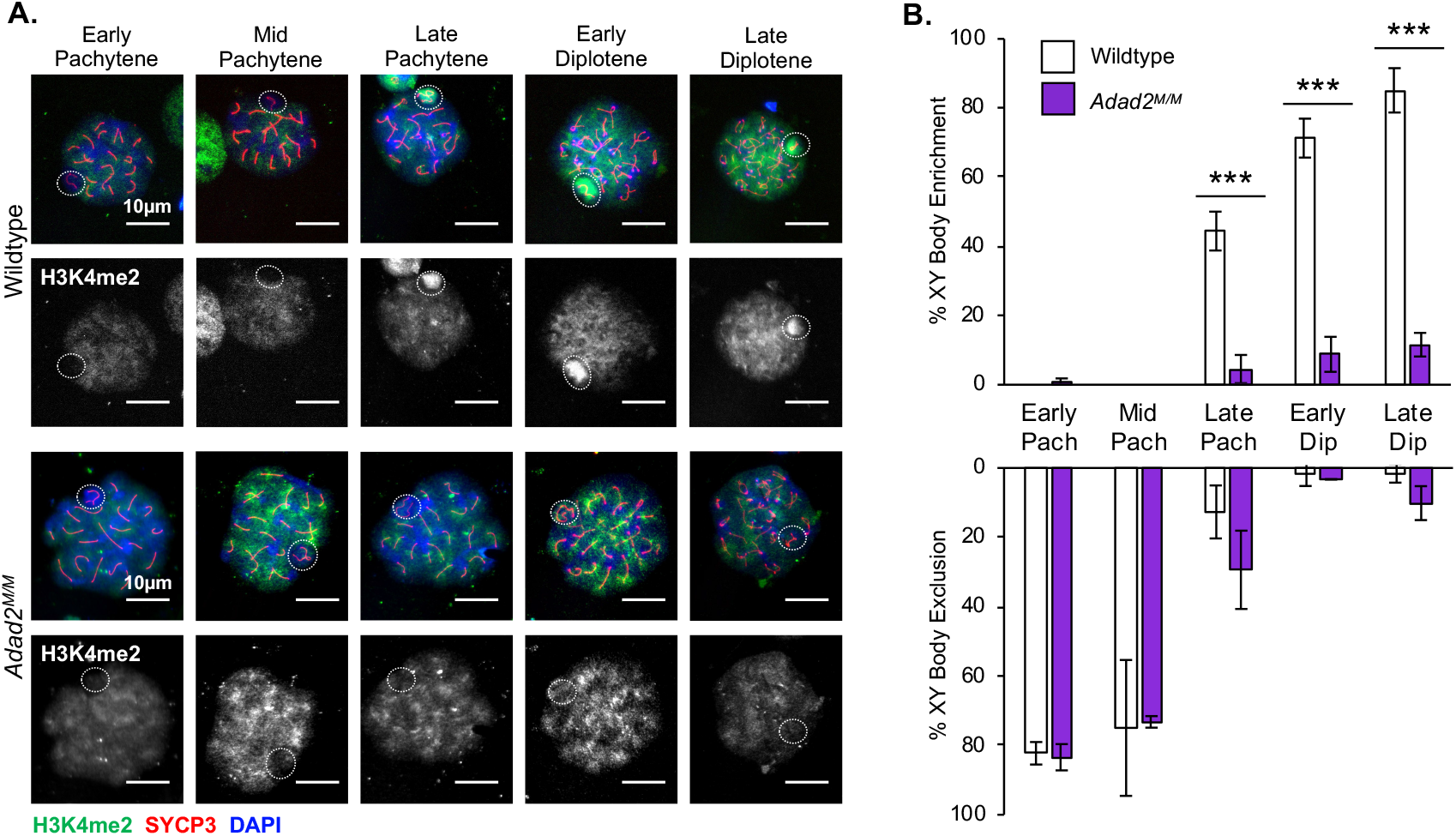
*Adad2* mutant spermatids do not form PMSC and fail to undergo the histone to transition protein transition. **A)** Immunofluorescence of H3K27me3 and H3K9me3 in adult wildtype and *Adad2^M/M^* round spermatids showing mutant round spermatids lack an H3K9me3-only chromatin domain (green - H3K27me3, red - H3K9me3, blue - DAPI. 63X magnification. Box – region of inset). **B)** Immunofluorescence of H3K27me3 and H3K4me2 in adult wildtype and *Adad2^M/M^* round spermatids demonstrating lack of H3K4me2 enrichment or exclusion in mutant round spermatids (green - H3K27me3, red - H3K4me2, blue - DAPI. 63x magnification. Box – region of inset). **C)** Immunofluorescence of TNP1 (green – TNP1, red – SYCP3, blue – DAPI) in adult wildtype and mutant testis sections by stage. TNP1 is imported into the nucleus starting in stage XI in wildtype spermatids, properly associates with the DAPI-dense regions of elongating spermatids by stage I, and signal begins to dissipate by stage IV. These events are delayed or abnormal in mutant elongating spermatids.

Our earlier TUNEL assays demonstrated a significant increase in round spermatid apoptosis and a general reduction of round spermatids. In spite of this and a lack of epididymal sperm (Snyder et al., 2020), occasional morphologically abnormal elongating spermatids are observed in *Adad2* mutant testes.

These cells provided an opportunity to examine post-meiotic chromatin remodeling in conditions of abnormal meiotic chromatin. To do so we examined the localization of the histones replacing protein transition protein 1 (TNP1) (Fig. 7C). In wildtype spermatids, the process of elongation accompanies dramatic reorganization of chromosome centromeres, marked by PCH, along the axis of elongation (Meyer-Ficca et al., 1998). Centromere elongation can be visualized by the DNA staining dye DAPI as spermatids mature. In wildtype elongating spermatids, the single chromocenter (region of intense DAPI staining) gives rise to an elongated region, the formation of which coincides with nuclear import of TNP1. *Adad2* mutant spermatids fail to undergo this chromocenter elongation, even those cells containing only a single chromocenter. Likewise, wildtype spermatids generate TNP1 and import it into the nucleus, where it roughly co-localizes with regions of DAPI in a stage dependent manner. However, in *Adad2* mutant spermatids TNP1 nuclear import is delayed and once imported, TNP1 shows poor co-localization with DAPI staining. Together, these observations suggest that the aberrations in epigenetic marks, and subsequent heterochromatin structures, of mutant meiotic germ cells directly impede their ability to undergo the histone to transition protein transition and the concurrent morphological elongation necessary to generate mature sperm.

## Discussion

Heterochromatin remodeling is a hallmark of germ cell differentiation. Herein we show loss of ADAD2, an RNA binding protein observed exclusively in mid-to late pachytene spermatocytes, leads to abnormal meiotic and post-meiotic heterochromatin in both the autosomes and sex chromosomes. This outcome is likely driven by a reduction late in meiosis of the DNA damage response protein MDC1 (Fig. 8). MDC1 is expressed throughout male germ cell meiosis and has previously been associated with establishment of the unique XY-body epigenetic state early in pachynema (Broering et al., 2014, Ichijima et al., 2011), though its role later in meiosis has been enigmatic. In *Adad2* mutants, MDC1-dependent XY-body epigenetic marks (Alavattam et al., 2016, Sin et al., 2012, Adams et al., 2018) are established properly but fail to be maintained through late meiosis. Additionally, γH2AX and ATR are aberrantly observed on autosomal axes late in meiosis in mutant spermatocytes. Although *Adad2* mutant spermatocytes complete meiosis, autosomal and sex chromosome heterochromatin is severely perturbed in post-meiotic spermatids resulting in high rates of post-meiotic spermatid apoptosis and failed post-meiotic chromatin compaction. These results explain *Adad2* mutants’ loss of post-meiotic germ cells, which are temporally far removed from ADAD2 expression, and additionally suggest *Adad2* mutants may be a unique model in which to query the function of MDC1 late in meiosis.

**Figure 8:**
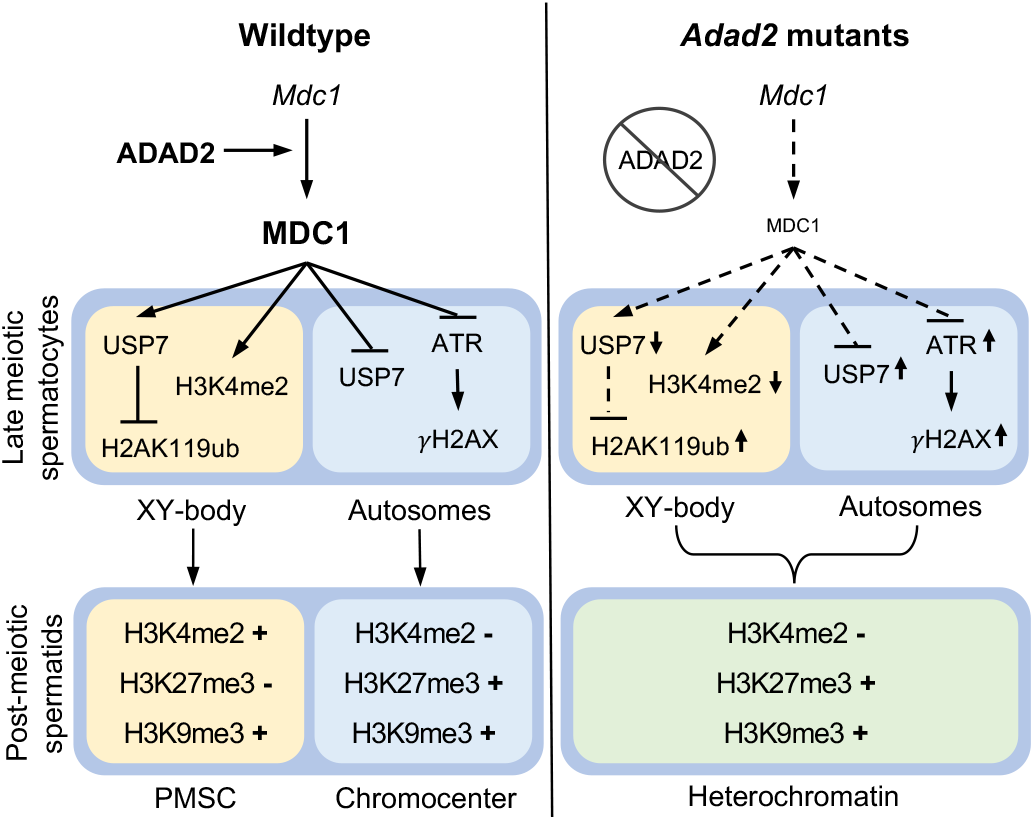
Summary of ADAD2 action and drivers of the *Adad2* mutant phenotype. In wildtype germ cells ADAD2 facilitates translation of *Mdc1* late in meiosis thus ensuring the correct epigenetic state of the XY body and autosomes and resulting in two unique heterochromatin domains (PMSC and the chromocenter) in the subsequent post-meiotic germ cells. In the absence of ADAD2, MDC1 is low late in meiosis leading to abnormal XY-body and autosome epigenetic signatures in late meiotic spermatocytes. In mutant post-meiotic spermatids, improperly marked chromatin domains result in the loss of distinct chromatin compartments.

The loss of MDC1 in *Adad2* mutants appears at first glance to be counterintuitive as the *Mdc1* transcript shows dramatically increased ribosome association, normally associated with increased translation. However, several other transcripts also demonstrate this unusual signature wherein overall transcript abundance is minimally decreased, ribosome association increased, and protein abundance dramatically reduced. Recent evidence from other systems has shown this phenomenon is indicative of ribosome dysfunction, specifically ribosome collisions arising from abnormal translation elongation dynamics (Simms et al., 2017, Yan and Zaher, 2021). Translation elongation is increasingly being recognized as an important aspect of translation control that can have a profound influence on cell physiology. Elongation rates are exceptionally dynamic during meiosis in yeast (Sabi and Tuller, 2019) and profoundly influence protein production during early embryogenesis (Richter and Coller, 2015). Further, evidence from mitotic models has demonstrated elongation can be regulated via post-translational regulation of the core elongation complex (Sivan et al., 2011). In spite of this, translation elongation is almost entirely unstudied in the male germ cell. The dramatic reduction of a key elongation factor, eEF1G, in *Adad2* mutants makes it tempting to postulate ADAD2 may be required for normal translation elongation in meiotic cells. The exact mechanisms by which ADAD2 may do so and how this relates to ADAD2’s distinct granular localization are, as yet, unknown. Ongoing and future efforts are aimed at answering these exciting questions.

The opposing impact of ADAD2 loss on BRCA1 and MDC1 provides a platform from which to dissect the relative contributions of both late in meiosis. In *Adad2* mutants, BRCA1 is dramatically up-regulated late in meiosis but, in spite of both axial ATR and γH2AX on the autosomes, BRCA1 is not observed outside of the XY-body. These findings support the notion that while BRCA1 acts as a nucleating factor in the XY-body for MDC1 recognition of γH2AX early in meiosis (Turner et al., 2004, Broering et al., 2014, Scully et al., 1997), thereafter BRCA1 alone is not sufficient to maintain sequestration of ATR to the XY-body. Further, BRCA1 is not required for ATR to associate with chromosome axes outside of the XY-body as autosomal ATR is observed in *Adad2* mutant spermatocytes in the absence of autosomal BRCA1. Future studies will be required to determine whether BRCA1 is dispensable late in meiosis for ATR maintenance in the XY-body.

Deregulation of MDC1-mediated events explains a large portion of the *Adad2* phenotype and provides novel insight into the function of MDC1 late in meiosis. Two MDC1-influenced epigenetic marks, H2K119Ub and H3K4me2 (Maezawa et al., 2018, Adams et al., 2018, Sin et al., 2012), have abnormal abundance and/or shifts from their normal autosomal or sex chromatin domains in *Adad2* mutant cells. As both have been associated with MSCI and post-meiotic X activation (Hasegawa et al., 2015), their alterations are a potential mechanism leading to the subtle X-chromosome expression defects observed in *Adad2* mutants. Additionally, MDC1 prevents H3K27me3 localization to the X and Y during meiosis (Ichijima et al., 2011), ultimately giving rise to the H3K27me3-negative PMSC in post-meiotic germ cells. Lack of detectable PMSC in *Adad2* mutants demonstrates failed H3K27me3 exclusion from the X and Y during meiosis. Together, these findings demonstrate MDC1 is required in mid-to late meiosis to establish specific epigenetic compartments in the autosomes and sex chromosomes of male germ cells.

Previous work has proposed MDC1 facilitates establishment of the γH2AX domain on the sex chromosomes via tethering of ATR and TOPBP1 (Abe et al., 2020, Ichijima et al., 2011). Redistribution of ATR and γH2AX late in meiosis in *Adad2* mutant spermatocytes demonstrates MDC1 tethering is required throughout the remainder of meiosis as well, while increased total γH2AX suggests MDC1 may also suppress ATR activity. This is further supported by the observation of increased H3K9me3 in *Adad2* mutant testes as the ATR/γH2AX network is upstream of SETDB1, which deposits H3K9me3 (Hirota et al., 2018). Future studies will be required in order to determine how ATR recognizes autosome axes when XY-body MDC1 is low.

Meiosis is not the only context in which MDC1 appears to tether components to regions containing γH2AX. During mitosis, DNA repair and DNA damage checkpoints are suppressed in order to ensure chromosome stability (Rieder and Cole, 1998). As such, mitotic cells rely on MDC1 to bind γH2AX at sites of DNA damage and mark them for repair during the next G1 phase (Orthwein et al., 2014, Lee, 2014). Recent work has shown MDC1 recruits and tethers TOPBP1 in mitosis to sites of DNA damage, thus forming a repair poised complex (Leimbacher et al., 2019). Meiotic cells represent another system in which DNA repair is suppressed (Srivastava and Raman, 2007) and, based on this work, also rely on the tethering function of MDC1 throughout cell division. As the composition of the mitotic DNA bound MDC1 complex remains undefined, it seems feasible MDC1 may also mediate epigenetic remodeling in mitosis at sites of DNA damage.

This work demonstrates ADAD2 is required to maintain a proper epigenetic state late in meiosis. Importantly, abnormalities in epigenetic marks are retained in mutant germ cells from late meiosis into post-meiotic spermatids, which show dramatic alterations in both autosome and sex chromosome heterochromatin. Only a handful of mutant models display similar post-meiotic germ cell heterochromatin defects as those observed in *Adad2* mutants (Maezawa et al., 2018, Shang et al., 2007, Martianov et al., 2002). In each, the causative events initiate late in meiosis and lead to substantial post-meiotic germ cell differentiation defects. These, and the findings reported here, demonstrate post-meiotic germ cell differentiation is highly reliant on proper heterochromatin remodeling late in meiosis. Further, they suggest meiotic germ cells lack a heterochromatin-dependent checkpoint late in meiosis, identifying mid-to late pachytene as a period in germ cell development that may be especially sensitive to the influence of epigenetic modifiers.

Overall, our findings define the mechanism whereby ADAD2 expression in meiotic germ cells leads to post-meiotic germ cell loss and reveal a heretofore unappreciated role for MDC1 in the maintenance of XY-body epigenetic state late in meiosis, underscoring the importance of MDC1 throughout meiosis and beyond. Lastly, we observed an unanticipated mode of translation regulation by ADAD2. The subsequent new avenues of study should shed light on an understudied level of translation regulation in meiotic germ cells and its connection to heterochromatin and the cell cycle.

## Materials and Methods

### Animal care and model generation

All animal use protocols were approved by the Rutgers University animal care and use committees. Mouse procedures were conducted according to relevant national and international guidelines (AALAC and IACUC). Generation of *Adad2^M/M^* mice was as described in (Snyder et al., 2020). RiboTag mice (Sanz et al., 2009) were obtained from The Jackson Laboratory and *Adad2-Ribotag* mice generated as in (Chukrallah et al., 2020). Mice were housed in a sterile, climate-controlled facility on a 12-hour light cycle. Mice were fed LabDiet 5058 irradiated rodent chow and had access to food and water *ad libitum*.

### Preparation of meiotic spreads and immunocytochemistry

Meiotic spreads were prepared following a modified version of the protocol described in (La Salle et al., 2008). In brief, testes from 30 dpp *Adad2^M/M^* and wildtype male mice were collected in 1x PBS and lysed in a hypotonic extraction buffer (30 mM Tris pH 8.2, 50 mM sucrose, 17 mM sodium citrate, 5 mM EDTA, 2.5 mM DTT and 0.5 mM PMSF) for fifteen minutes at room temperature. Tubules were transferred to and dispersed in a 100 mM sucrose solution. Twenty μL of tubule solution was spread on charged slides coated with 1% PFA solution plus 0.14% Triton X-100 and left in a humid chamber to dry overnight. Slides were washed for 2 minutes in 0.4% Kodak Photoflo and air-dried at room temperature before either using directly for immunocytochemistry or stored at −20°C for later use.

Prior to staining, slides were blocked for one hour at room temperature in 1X ADB (45 mM BSA, 1% normal goat serum, and 0.2% Triton X-100 in 1x PBS). Primary antibodies were diluted in 1X ADB and incubated overnight in a humid chamber either at 4°C or room temperature. Slides were then rinsed in 1X PBS for 10 minutes, then twice in 1X ADB for 10 minutes each. Fluorescent secondary antibodies were diluted in 1X ADB. Slides were incubated in light-protected humid chamber for 2 hours at room temperature, then washed in 1X PBS for 10 minutes three times. Slides were mounted using DAPI Fluoromount-G (SouthernBiotech) and stored at 4°C with light protection. For a detailed breakdown of antibody concentrations and conditions, please see Supplemental Table 3.

Cells were imaged using a custom-built Zeiss microscope with brightfield and fluorescent capabilities. Each channel was imaged individually through MetaMorph imaging software (Molecular Devices) and color-combined using the program’s built-in color combine tool. Provided images representative of biological triplicate or greater. With the exception of RPA2, all quantification was done via direct visualization. A minimum of thirty cells per meiotic stage per sample were quantified. For all quantifications, slides from a total of three wildtype and three *Adad2^M/M^* samples were used. Spermatocyte stages were distinguished by SYCP3 pattern as in (Luo et al., 2015). Briefly, X and Y chromosome structure was used to distinguish early, mid, and late pachytene spermatocytes while desynapsis of three or fewer SYCP3 strands indicated early diplotene and greater than three late diplotene.

### RPA2 foci quantification

Thirty spermatocytes at each stage (early, mid, and late pachytene as well as diplotene) were imaged per sample (30 dpp wildtype and *Adad2^M/M^* n = 3). For each, an RPA2 image, an SYCP3 image, and DAPI image, were taken. For each cell, RPA2 foci were quantified using ImageJ (Abramoff04) and the Analyze Particle function (minimum size = 10 pixels; threshold = 50). Cell boundary was determined by DAPI signal and stage by SYCP3 as above. The number of discrete foci was recorded and averaged by spermatocyte stage and genotype. Data was visualized in R.

### Protein isolation and Western blotting

Testes were collected from adult (60-70 dpp) and 21 dpp *Adad2^M/M^* and wildtype male mice and flash frozen. Tissue was ground on liquid nitrogen and total protein extracted by RIPA buffer with protease inhibitors at a ratio of 1 mL buffer to 100 mg tissue. Protein concentration was determined via the DC protein assay (BioRad) as described by (Snyder et al., 2020). Twenty μg protein was electrophoresed on 10% acrylamide gels per sample and Coomassie used to verify equal protein loading. Following wet transfer of proteins to PVDF membrane (BioRad), membranes were blocked and incubated overnight with primary antibody at 4°C. Images were developed with SuperSignal West Pico PLUS Chemiluminescent Substrate (Thermo Scientific) and visualized using an Azure Biosystems C600 imager.

Western blot band intensity was quantified using ImageJ as described in (Janes, 2015). A band intensity value was calculated for each sample in the blot using ImageJ’s Analyze Gel tool. The band density of the experimental blot was normalized by taking the ratio of the density of the experimental band to the density of the corresponding GAPDH or Total H3 (histone extraction only) band. Wildtype and *Adad2^M/M^* sample values were averaged. Standard deviation was calculated, and significant differences determined by Student’s t-test.

### Extraction and Western Blotting of Histones

Testes were collected from adult (60-70 dpp) and 21 dpp *Adad2^M/M^* and wildtype male mice, ground on dry ice, and resuspended in Triton extraction buffer (TEB) (0.5% Triton X-100, 2 mM phenylmethylsulfonyl fluoride (PMSF), 0.02% NaN3) at 100 mg tissue/mL buffer. Tissue was lysed by rotating at 4°C for 10 minutes, then centrifuged at 2000xG for 10 minutes at 4°C. Supernatant was discarded, and pellet was resuspended in half the starting volume of TEB and centrifuged as above. Pellet was resuspended in an equal volume of 0.2N HCl and acid-extracted overnight at 4°C. Sample was then centrifuged at 2000 x g for 10 minutes at 4°C and supernatant collected. Protein content was determined using the BioRad DC protein assay, as above, and equal loading confirmed by Coomassie. Twenty μg histone lysate was electrophoresed on 15% acrylamide gels and transferred to PVDF membranes as above. Membranes were washed in 1X PBS with 0.1% Triton and blocked in 5% BSA in 1xPBS with 0.1% Triton. Membranes were visualized and imaged as above.

### Histological evaluation

Adult and 21 dpp wildtype and *Adad2^M/M^* testes were collected and fixed overnight in Bouin’s Solution (Sigma Aldrich). Tissue was cleared in diH_2_O, dehydrated in increasing concentrations of ethanol, paraffin embedded, and cut to 4 μm sections. Slides were deparaffinized and rehydrated before staining with Periodic Acid (Millipore Sigma), Schiff’s Reagent (Millipore Sigma), and Meyer’s Hematoxylin (Millipore Sigma). Slides were dehydrated in ethanol and mounted with Permount mounting medium (Fisher Scientific).

Histological parameters as laid out in (Lonnie D. Russel, 1990) were used to 1) quantify the number of round spermatids per tubule and number of round spermatid-containing tubules per 21 dpp sample and 2) the number of MII spermatocytes (n > 100 / sample) per tubule and the number of stage XII tubules (n = 50) per adult sample. For both counts, totals and averages for each genotype were calculated, as well as standard deviation. Student’s *t-*test was used to identify significant differences by genotype.

Round spermatid morphology was quantified in DAPI-stained samples (21 dpp wildtype and *Adad2^M/M^*, n = 3). At least 1200 round spermatids were quantified per sample. Spermatids were binned by number of intensely DAPI-staining structures (1, 2, or 3+). Counts were averaged by genotype percent of each pattern was calculated. Standard deviation was calculated, and significant differences determined by Student’s t-test.

### Immunofluorescence

Testes were dissected from adult mice and fixed overnight in 4% PFA. Tissue was rinsed in PBS and dehydrated in increasing concentrations of ethanol before embedding in paraffin wax. Four μm sections were utilized for all applications. Antigen retrieval was performed by boiling slides in either 14.3 mM citrate solution (pH 5.95) for 7 minutes, 10 mM glycine (pH 2.5) for 8 minutes, or Tris-EDTA (10 mM Tris-HCl, 1 mM EDTA, 0.05% Tween; pH 9.0) for thirty minutes. Slides were mounted using DAPI Fluoromount-G and stored at 4°C with light protection. Slides were visualized on a custom-built microscope (Zeiss) with fluorescent and brightfield capabilities. Provided images representative of biological triplicate or greater. Signal intensity was matched across slides by matching background (interstitial) signal intensity Developmental stages were determined according to the parameters set forth in (Lonnie D. Russel, 1990), facilitated by SYCP3 co-staining where possible.

### TUNEL Assay

Adult testes from wildtype and *Adad2^M/M^* mice were PFA-fixed, paraffin embedded, and sectioned as above. Slides were deparaffinized with xylenes and rehydrated in decreasing concentrations of ethanol (100%, 95%, 70%, and 50%) before incubation at 37°C in Proteinase K (1:500 in 1X PBS). TMR Red In Situ cell death detection Kit (Millipore Sigma) was utilized, as per manufacturer’s instructions. Slides were mounted using DAPI Fluoromount-G and stored at 4°C with light protection. TUNEL positive cells were quantified by direct visualization. Cells determined via DAPI staining to be either spermatocytes or spermatids were quantified, as were the number of these cells with a distinct red TUNEL signal in the triple channel. A minimum of 100 cells per each type were quantified per sample (n = 3 for wildtype and *Adad2^M/M^*). From this, percent TUNEL-positive cells were calculated for each cell type in each sample, then averaged by genotype. Standard deviation was calculated, and significant differences determined by Student’s t-test.

### Ribotag RNA immunoprecipitation

Testes were collected from 21 dpp males heterozygous for *Rpl22-HA* and *Stra8 iCre* that were either *Adad2^M/M^* (n = 4) or *Adad2^+/+^* (n = 3). Immunoprecipitation and RNA extraction were carried out as described in (Chukrallah et al., 2020). In brief, testes were homogenized in lysis buffer, precleared twice using antibody-free beads, and pre-immunoprecipitation (input) fractions collected. Overnight incubation with an anti-HA antibody (ABCAM) was followed by two-hour incubation with Protein A Dynabeads (Invitrogen). Beads were washed and RNA was extracted using the miRNeasy mini kit (Qiagen) per manufacturer’s instructions. Each biological replicate generated a total RNA sample (input) and an immunoprecipitated RNA sample (IP).

### RNA Sequencing and Quality Control

Input and IP samples were quantified via Nanodrop and assessed for RNA integrity using an Agilent Bioanalyzer (Agilent RNA 6000 Pico). Sample RNA was sent to Genewiz (South Plainfield, NJ) for commercial sequencing (Total RNA, Illumina HiSeq 4000, paired end, 150bp reads). Strand specific libraries were prepped with the Ribo-zero Gold HMR and TruSeq ^®^ Stranded Total RNA Library Prep Human/Mouse/Rat.

Following sequencing, read quality was assessed using the FastQC software from Babraham Bioinformatics and summarized with MultiQC (V1.9, (Ewels et al., 2016)). *In silico* rRNA depletion was carried out using BOWTIE2 (V2.4.1, (Langmead and Salzberg, 2012)). Based on initial quality reports, the first 20 and last 30 base pairs were trimmed (TRIMMOMATIC, (Bolger et al., 2014)) generating 100 bp, paired end reads. A final rerun of FastQC and MultiQC indicated improved quality. Principle component analyses to confirm sample ID assignment were carried out using pcaExplorer (Marini and Binder 2019).

### Data analyses

An expanded testis transcriptome was generated by appending novel testis-specific genes and isoforms (Gamble et al., 2020) to the current Ensembl mouse transcriptome (Mus_musculus.GRCm38.90, mm10). Reads were aligned to this expanded transcriptome and abundance estimated using RSEM (v 1.3.3) (Li and Dewey, 2011). For input samples, differential expression was calculated by EBSeq (release 3.12) (Seirup et al., 2020) with a cutoff of PPDE ? 0.95 and a directional fold change greater than one. Ribosomal association (RA) was determined by calculating the ratio of IP TPM over input TPM. Any transcripts with any input values equaling zero were removed. A Welch t-test was conducted using R (matrixTests) to identify differential ribosome association (DRA, p < 0.05). Graphical representations of data as scatter plots, heat maps (base R), and violin plots (ggplot2) were generated using R. Ontological analyses were performed using DAVID bioinformatics database (Huang da et al., 2009). Ontological categories were identified from gene lists differentially expressed or differentially ribosome associated in wildtype or *Adad2^M/M^* samples. Clusters were identified using a medium classification stringency and p values were confirmed to be statistically significant (p < 0.05).

## Author Contributions

Lauren Chukrallah was the lead for experimental investigation, data visualization, and validation and supported methodology development. LC also wrote the original manuscript draft and supported review and editing. Aditi Badrinath provided data curation and formal analysis as well as supported data visualization. Gabrielle Vittor provided support for experimental investigation and validation. Elizabeth M. Snyder conceptualized the research idea, acquired funding, administered the project, supervised all involved authors, and was the lead for methodology and model development and review and editing of the manuscript.

## Acknowledgements

The authors would like to thank Kelly Seltzer, Christopher Eddy, and Gabriella Acoury for support with animal husbandry and colony management; Dr. William Belden for kindly providing the H3K27me3 antibody; and Drs. Tracy Anthony, Karen Schindler, Devanshi Jain, and Kim McKim for critical evaluation and intellectual contribution throughout project development. The authors would like to especially thank our funding sources: the Eunice Kennedy Shriver National Institute of Child Health and Human Development (NIH-NICHD F32 HD072628 and K99/R00 HD083521 to ES) and Rutgers University (to ES).

